# Selective agonism of GPR34 stimulates microglial uptake and clearance of amyloid β fibrils

**DOI:** 10.1101/2024.05.08.593262

**Authors:** Hayato Etani, Sho Takatori, Wenbo Wang, Jumpei Omi, Aika Akahori, Hirotaka Watanabe, Iki Sonn, Hideyuki Okano, Norikazu Hara, Mai Hasegawa, Akinori Miyashita, Masataka Kikuchi, Takeshi Ikeuchi, Maho Morishima, Yuko Saito, Shigeo Murayama, Takashi Saito, Takaomi C Saido, Toshiyuki Takai, Tomohiko Ohwada, Junken Aoki, Taisuke Tomita

**Affiliations:** Laboratory of Neuropathology and Neuroscience, Graduate School of Pharmaceutical Sciences, The University of Tokyo, 7-3-1 Hongo, Bunkyo-ku, Tokyo 113-0033, Japan; Department of Health Chemistry, Graduate School of Pharmaceutical Sciences, The University of Tokyo, Tokyo 113-0033, Japan; Department of Physiology, Keio University School of Medicine, Tokyo, 160-8582, Japan; Keio University Regenerative Medicine Research Center, Kanagawa, 210-0821, Japan; Department of Molecular Genetics, Brain Research Institute, Niigata University, Niigata, Japan; Department of Neuropathology (the Brain Bank for Aging Research), Tokyo Metropolitan Institute for Geriatrics and Gerontology, Tokyo 173-0015, Japan; Department of Neurocognitive Science, Institute of Brain Science, Nagoya City University Graduate School of Medical Science, 1 Kawasumi, Mizuho-cho, Mizuho-ku, Nagoya, Aichi 467-8601, Japan; Laboratory for Proteolytic Neuroscience, RIKEN Center for Brain Science, 2-1 Hirosawa, Wako, Saitama 351-0198, Japan; Department of Experimental Immunology, Institute of Development, Aging and Cancer, Tohoku University, 4-1 Seiryo, Sendai 980-8575, Japan; Department of Organic and Medicinal Chemistry, Graduate School of Pharmaceutical Sciences, The University of Tokyo, Tokyo 113-0033, Japan

**Keywords:** Alzheimer disease, Amyloid-β, Microglia, GPR34

## Abstract

Microglia, the primary immune cells of the central nervous system, play a crucial role in maintaining brain homeostasis through phagocytosis of various substrates, including amyloid-β (Aβ) fibrils, a hallmark of Alzheimer disease (AD) pathology. However, the molecular mechanisms regulating microglial Aβ uptake remain poorly understood. Here, we identified GPR34, a Gi/o-coupled receptor highly expressed in microglia, as a novel regulator of fibrillar Aβ phagocytosis. Treatment with a selective GPR34 agonist, M1, specifically enhanced uptake of Aβ fibrils, but not its monomer or oligomer, in both mouse and human microglia. Mechanistically, M1 reduced intracellular cAMP levels, which inversely correlated with Aβ uptake activity. Importantly, a single intrahippocampal injection of M1 in an AD mouse model significantly increased microglial Aβ uptake in vivo. Furthermore, single-nucleus RNA-sequencing analysis of Japanese AD patient samples revealed a significant reduction of GPR34 expression in microglia from AD patients compared to controls. We also observed an age-dependent decline in microglial GPR34 expression in both human and mouse datasets, suggesting a potential contribution of GPR34 downregulation to age-related Aβ accumulation and AD risk. Collectively, our findings identify GPR34 as a promising target for modulating microglial Aβ clearance and highlight the therapeutic potential of GPR34 agonists in AD.

**Significance statement:** Alzheimer disease (AD) is characterized by amyloid-β (Aβ) accumulation in the brain. Microglia, the brain’s immune cells, play a crucial role in the metabolism of Aβ. We discovered that activating the microglial receptor GPR34 with a selective agonist enhances the phagocytosis of Aβ fibrils, a key pathogenic form of Aβ. Importantly, GPR34 expression decreases with aging and AD progression, potentially contributing to impaired Aβ clearance. Our findings highlight GPR34 as a promising therapeutic target for AD, as boosting its activity could promote Aβ clearance and slow disease progression. This study provides valuable insights into microglial function in AD and offers a novel strategy for developing disease-modifying therapies.

## Introduction

Imbalances in the production and clearance of amyloid-β (Aβ) peptides in the brain are central to the pathogenesis of Alzheimer disease (AD). For example, mutations elevating Aβ production or aggregation are implicated in familial AD (1). Conversely, the *APP* A673T variant, which is associated with lower Aβ levels, significantly reduces the risk of late-onset AD (2). In addition, recently approved anti-Aβ antibodies have shown efficacy in slowing cognitive decline and disease progression in AD patients by improving Aβ clearance (3, 4). Taken together, these findings position cerebral Aβ levels as a critical determinant of AD pathogenesis and progression, and highlight drugs that promote Aβ clearance as potential preventive and disease-modifying therapies. However, the precise mechanisms underlying Aβ clearance remain to be elucidated.

As resident immune cells of the central nervous system (CNS), microglia phagocytose and degrade extracellular Aβ species. Recent genetic analyses have identified numerous genetic risk factors for AD, many of which are found in genes that are highly or specifically expressed in microglia (5, 6). Among these genes, *TREM2* encodes a microglial Aβ receptor, and its rare hypomorphic variant significantly increases AD risk (7, 8). *Trem2* deficiency in AD mouse models exacerbates Aβ deposition (9–11), highlighting the pathologically important role of TREM2-mediated Aβ clearance. Thus, impairments in microglial clearance pathways may promote AD pathogenesis, whereas enhancing microglial phagocytic activity holds promise for preventing or slowing disease progression. Therapeutics aimed at activating Aβ clearance by microglia may be beneficial in AD.

G-protein coupled receptor 34 (GPR34) is a Gi/o-coupled receptor highly expressed in mononuclear cells of the immune system. In the CNS, GPR34 is specifically expressed in microglia, especially in the homeostatic state (12, 13). Together with P2RY10 and GPR174, GPR34 comprises the lysophosphatidylserine (LysoPS) receptor family (14, 15). Uniquely within this receptor family, GPR34 exhibits selectivity for LysoPS species with a fatty acid at the *sn*-2 position (16–19). While GPR34 is involved in various immunomodulatory functions (20–22), another study highlights its importance in the regulation of phagocytosis in microglia (23). Regarding pathological associations, GPR34 is involved in the neuropathic pain after nerve injury (24). However, the relationship between GPR34 and AD pathology remains largely unknown, in particular whether GPR34 contributes to Aβ clearance by microglia.

In this study, we report that activating GPR34 with our recently developed selective agonist M1 (referred to as compound **14** in (18)) enhances microglial phagocytosis of Aβ. In mouse primary microglia, M1 specifically increased uptake of Aβ fibrils without affecting Aβ monomers or oligomers. The underlying mechanism involves the suppression of cyclic AMP (cAMP) levels in microglia. Enhancement of Aβ phagocytosis by M1 was also observed in human microglia-like cells and further validated *in vivo* using an AD mouse model. In addition, we found that expression of *GPR34* decreases with age and shows its reduction in human AD brains correlating with amyloid deposition. Taken together, these results demonstrate that GPR34 activation promotes microglial clearance of Aβ fibrils, suggesting that age-related GPR34 reductions may contribute to AD pathogenesis.

## Results

### Selective agonism of GPR34 specifically increases microglial uptake of Aβ fibrils

To investigate the effect of the GPR34 agonist M1 (Fig. 1*A*) on microglial uptake of Aβ, we adapted a flow cytometry-based assay previously described by Mandrekar *et al.* (25). This assay measures cellular fluorescence derived from fluorophore-conjugated, internalized Aβ fibrils (fAβ). Exposing murine primary microglia to fluorescent fAβ for 3 hours resulted in an elevated fluorescence profile, which typically appeared as a broad peak or, under certain experimental conditions, as two distinct peaks (Fig. 1*B*). The peak with higher fluorescence intensity corresponds to cells that have phagocytosed high-molecular-weight Aβ aggregates, consistent with the previous findings (25). Our analysis focused on both the proportion and the mean fluorescence intensity (MFI) of cells exhibiting this higher fluorescence. We found that treatment with M1 at concentrations above 10 nM led to an approximate 35% increase in the MFI of fAβ^+^ cells (Fig. 1 *B* and *C*), without affecting the proportion of fAβ^+^ cells (*SI Appendix*, Fig. S1 *A*–*C*). This suggests that M1 increases the quantity of fAβ engulfed by individual microglial cells. The positive effect of M1 was not observed at concentrations below 1 nM, consistent with the previously reported EC_50_ value of 5 nM for M1 (18). Furthermore, the effect was evident as early as 1.5 hours after exposure to M1 and persisted for at least 7.5 hours (Fig. 1*D*). To confirm that the observed effect is mediated through GPR34, we assessed the impact of M1 in *Gpr34* knockdown cells. Two independent siRNAs targeting *Gpr34* resulted in a substantial reduction in *Gpr34* expression (Fig. 1*E*). Notably, *Gpr34* knockdown itself led to a significant decrease or a trend towards reduced uptake of fAβ (Fig. 1*F*), suggesting that GPR34 positively regulates fAβ phagocytosis. Furthermore, *Gpr34* knockdown abrogated the M1-induced increase in fAβ uptake (Fig. 1*F*), indicating that GPR34 is indeed the target of M1.

**Fig.1:**
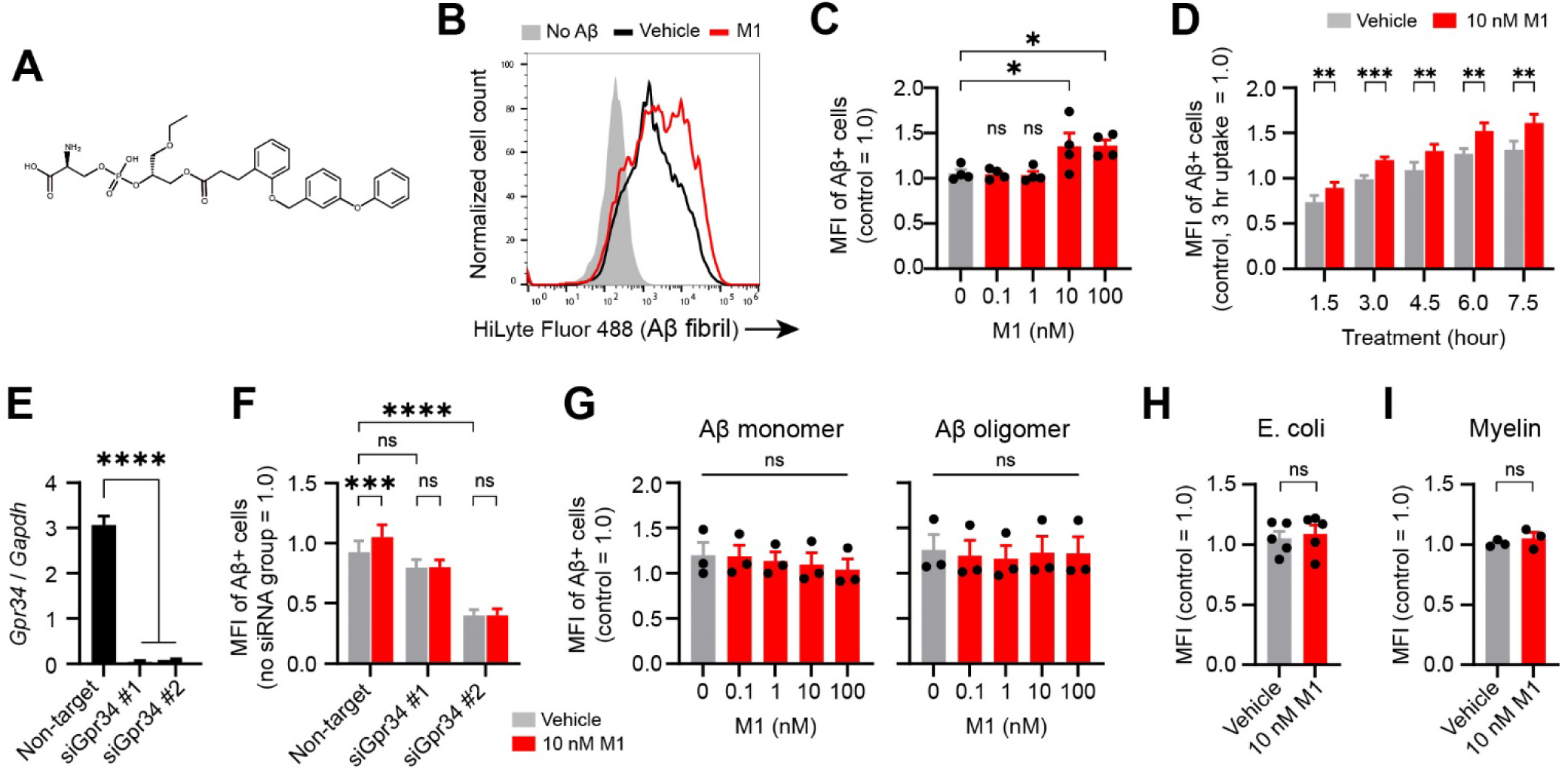
GPR34-selective agonist M1 specifically increased fibrillar Aβ uptake by murine primary microglia (***A***) The chemical structure of the GPR34 agonist M1. (***B***) Representative flow cytometry plots showing uptake of fluorescently labeled Aβ fibrils (fAβ) by mouse primary microglia pretreated with 10 nM M1 or vehicle, after 3-hours incubation with fAβ. (***C***) M1 increased the mean fluorescence intensity (MFI) of fAβ+ cells in a concentration-dependent manner after 3 hours of fAβ treatment. Data are normalized to the MFI of control cells treated with fAβ alone (*i.e.*, without M1 or vehicle treatment) (N=4; Dunnett’s test). (***D***) Time course analysis showing the effect of 10 nM M1 on fAβ uptake by microglia (N=7–8; paired t-test). (***E***) Knockdown efficiencies of two independent *Gpr34* siRNAs were validated by RT-qPCR (N=3, Dunnett’s test). (***F***) The M1-mediated increase in fAβ uptake was abolished by *Gpr34* knockdown using these siRNAs (N=5– 6; two-way repeated measures ANOVA followed by Šídák’s multiple comparisons test). ***G***–***I*** M1 did not affect microglial uptake of fluorescently labeled Aβ monomers or oligomers (***G***, N=3, Dunnett’s test), *E. coli* particles (***H***, N=5, paired t-test) or myelin debris (***I***, N=3, paired t-test). Bars represent the mean ± SEM.

Soluble Aβ species are taken up differently than fAβ, through non-phagocytic mechanisms such as macropinocytosis (25). Therefore, we examined the effects of M1 on the uptake of Aβ monomer and oligomer. At any concentrations, M1 did not significantly alter the MFI of fluorescence-positive cells for these soluble Aβ (Fig. 1*G*, *SI Appendix*, Fig. S1 *D* and *E*), suggesting that M1 has a specific enhancing role in fAβ phagocytosis. To further examine the specificity of M1-mediated phagocytosis enhancement, we conducted a similar experiment using two known phagocytosis substrates, *Escherichia coli* (*E. coli*) and myelin debris. Notably, M1 failed to enhance their uptake (Fig. 1 *H* and *I*). This finding indicates that M1 distinctly influences the uptake mechanism of fAβ.

### Mode of action of M1

GPR34 is a Gi/o-coupled GPCR that suppresses adenylyl cyclase and activates downstream kinases such as AKT and ERK upon ligand binding (Fig. 2*A*). Consistent with this, M1 treatment induced time-dependent increases in phosphorylated AKT and ERK levels (*SI Appendix*, Fig. S2). We also investigated whether M1 alters cAMP levels. To elevate the typically low steady-state cAMP levels, we pretreated cells with forskolin, an activator of adenylyl cyclase. We then observed an expected rise in intracellular cAMP concentration following forskolin treatment. However, when co-treated with M1, this elevation in cAMP levels was counteracted (Fig. 2*B*). Considering that an elevated cAMP level suppresses phagocytic activity (26–28), we investigated the effects on the fAβ uptake in these treated cells. We found that the single treatment of forskolin was enough to reduce the MFI of fAβ^+^ cells, while the co-treatment of M1 restored the impaired fAβ uptake back to the basal level (Fig. 2*C*). Interestingly, forskolin specifically affect the MFI without altering the proportion of fAβ^+^ cells like the effect of M1 (*SI Appendix*, Fig. S3). Moreover, forskolin treatment did not alter the uptake of monomeric and oligomeric Aβ species (Fig. 2 *D* and *E*). These data indicate that cAMP levels inversely correlate with the phagocytic activity specifically for fAβ, which likely underlies the substrate-specific effect of M1.

**Fig.2:**
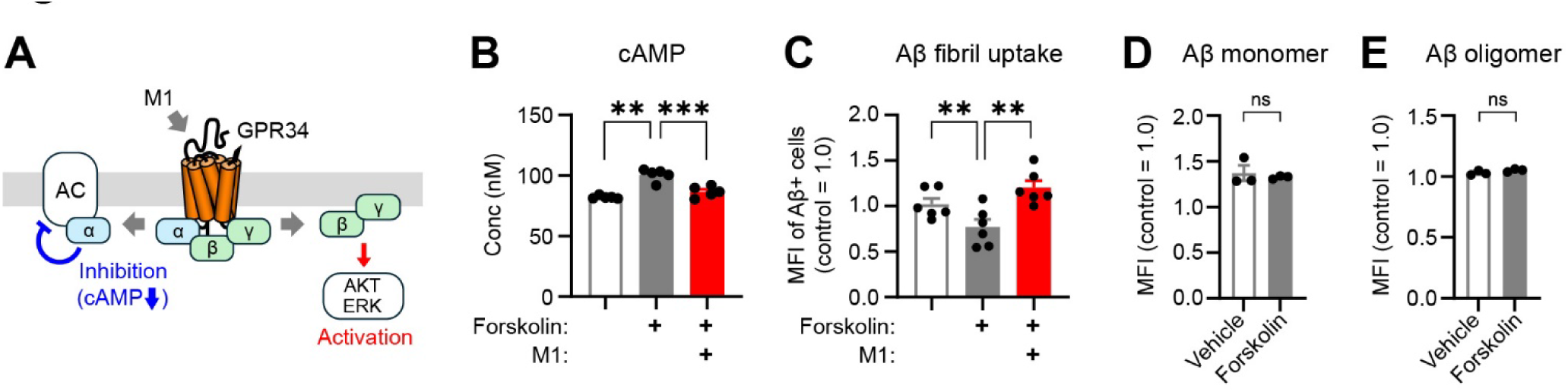
M1 reduces intracellular concentration of cAMP (***A***) Schematic illustrating the proposed mechanism by which GPR34 activation reduces intracellular cAMP levels. (***B***) Treatment with the adenylyl cyclase (AC) activator forskolin increased intracellular cAMP levels in primary microglia, which was counteracted by co-treatment with 10 nM M1 (N=5, one-way repeated measures ANOVA and Tukey’s test). ***C***–***E*** Forskolin treatment suppressed microglial uptake of fAβ. This impairment was rescued by co-treatment with M1 (***C***, N=6, one-way repeated measures ANOVA and Tukey’s test). In contrast, forskolin did not significantly affect microglial uptake of monomeric (***D***) or oligomeric (***E***) Aβ species (N=3, paired t-test). Bars represent the mean ± SEM.

### M1 promotes microglial uptake of Aβ in vivo

To explore the potential of M1 in promoting microglial uptake of Aβ *in vivo*, we utilized the mutant *APP* knock-in *App^NL-G-F^*mouse model (hereafter NLGF mice) (29). These mice, aged between 15–17 months, were administered a single injection of M1 into the hippocampus. The next day, brain Aβ deposits were fluorescently labeled by intraperitoneal injection of Methoxy-X04 (MX04). We then subjected these brains to flow cytometry analysis of the CD11B+/CD45+ microglial population to quantify MX04-positive cells (Fig. 3*A*). Our analysis revealed a notable increase in the MX04-positive microglial fraction upon M1 administration (Fig. 3 *B* and *C*). This observation indicates that M1 effectively enhances the microglial uptake of Aβ fibrils in an *in vivo* context.

**Fig.3:**
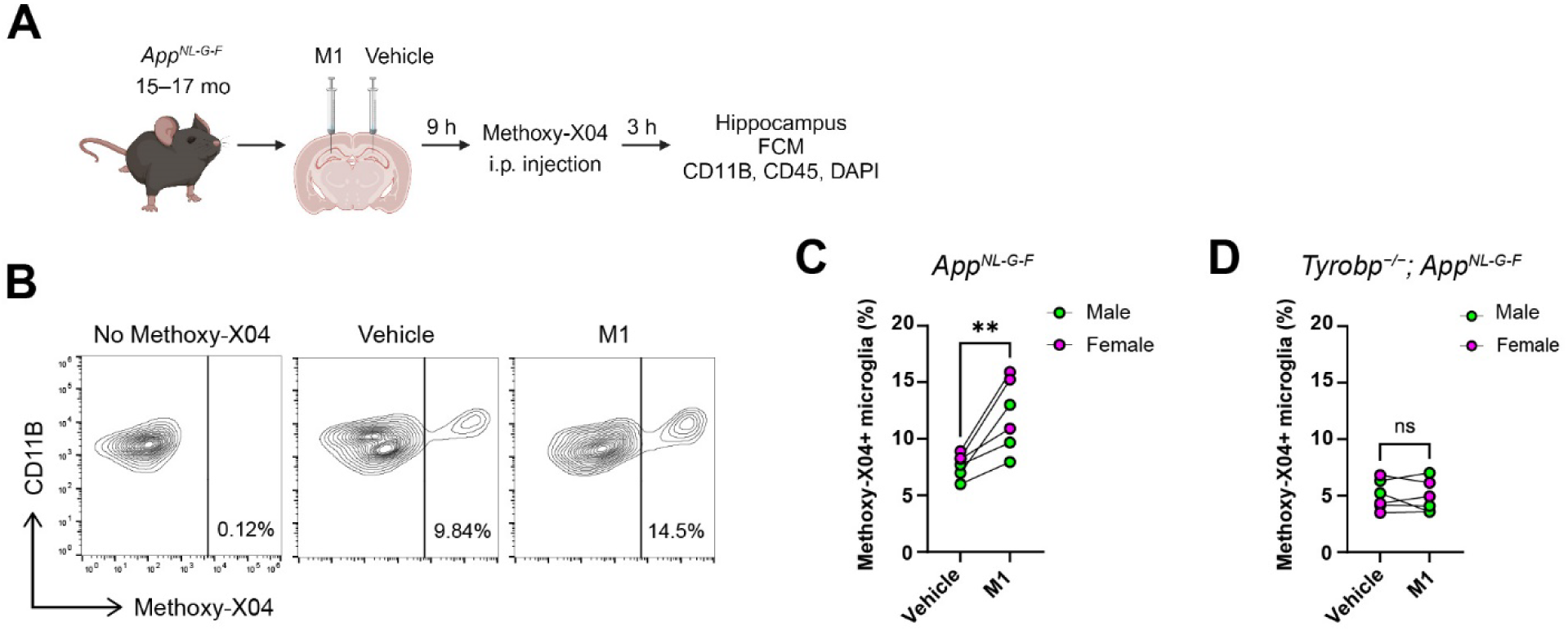
The effect of M1 on microglial uptake of Aβ fibrils *in vivo* (***A***) Experimental design schematic. 15–17 month-old male and female NLGF mice received bilateral intrahippocampal injections of vehicle or M1 (250 µM, 0.8 µL), followed by intraperitoneal injection of the Aβ tracer Methoxy-X04 (MX04; 10 mg/kg) the next day. Hippocampi were harvested 12 hours after M1 injection for flow cytometry analysis of MX04+ microglia (CD11B+/CD45+). ***B***, ***C*** Representative flow cytometry plots (***B***) and quantification (***C***) showing an increase in the percentage of MX04+ microglia upon M1 administration. (***D***) In *Tyrobp*-deficient NLGF mice lacking functional TREM2, M1 failed to increase the percentage of MX04+ microglia. Each pair of data points connected by a line represents a single experiment (N=6; male and female mice are indicated in *green* and *magenta*, respectively). Bars represent the mean ± SEM. Paired t-test.

Given that *Trem2* deficiency impairs Aβ metabolism in β-amyloidosis mice, this receptor molecule is important for microglial recognition and uptake of Aβ. Consequently, we asked whether the effect of M1 is somehow related to TREM2 function. For this purpose, we employed *Tyrobp*-deficient mice (30), which lack a functional TREM2 receptor. We noted a significant reduction of MX04-positive microglia in *Tyrobp*-deficient NLGF mice. Remarkably, M1 administration did not augment the MX04-positive microglial fraction in these mice (Fig. 3*D*). These data suggest that M1’s action requires the function of TREM2/TYROBP in microglia.

### M1 promotes fAβ uptake by human iPSC-derived microglia-like cells

Given the beneficial effect of M1 in promoting fAβ uptake by microglia, M1 emerges as a potential therapeutic candidate to boost fAβ metabolism in the brain. To ascertain whether M1 exerts a comparable impact on human-derived microglia, we employed a recently developed method for the efficient differentiation of microglia-like cells (iMGLs) from human induced pluripotent stem cells (iPSCs) (Fig. 4*A*). This method involves using iPSCs genetically modified to express the transcription factor *SPI1*, a crucial factor for microglial differentiation (31). Implementing this protocol, we were able to derive iMGLs exhibiting characteristic microglial projections (Fig. 4*B*). Subsequently, we subjected these human iMGLs to the flow-cytometric analysis to assess the effect of M1 on fAβ uptake (Fig. 4*C*). Notably, despite variations in the effective concentration range, the stimulatory effect of M1 on fAβ uptake in human cells mirrored what we observed in mouse primary microglia (Fig. 4*D*).

**Fig.4:**
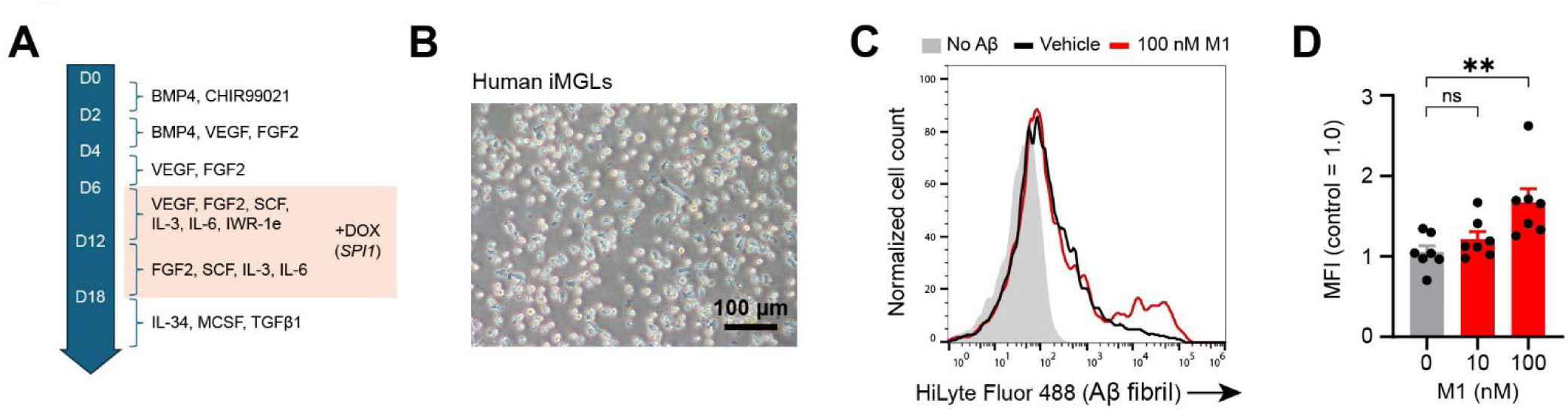
The effect of M1 in human iPSC-derived microglia-like cells (***A***) Protocol for differentiating human iPSC-derived microglia-like cells (iMGLs) using a doxycycline (DOX)-inducible SPI1 expression system. (***B***) A representative image of human iPSC-derived microglia-like cells (Scale bar: 100 µm). ***C***, ***D*** Representative flow cytometry plots (***C***) and quantification (***D***) showing an increase in fAβ uptake by microglia upon M1 treatment (N=7). Bars represent the mean ± SEM. Dunnett’s test.

### Reduction of GPR34 expression in brain microglia associated with Alzheimer disease and aging

To investigate potential changes in GPR34 expression associated with AD, we performed single-nucleus RNA-sequencing (snRNA-seq) on postmortem brains of 15 Japanese AD patients and 8 normal controls (*SI Appendix*, Table S1). We successfully identified a distinct microglial cluster based on established marker genes (Fig. 5*A*). *GPR34* expression was specifically enriched in this microglial population (Fig. 5*B*). Notably, when comparing *GPR34* levels between AD patients and controls across the annotated clusters, we observed a significant reduction in *GPR34* expression within the microglial cluster from AD patients (log_2_FC = −0.49, adjusted *p* = 0.00004; Fig. 5*C*). Among the differentially expressed genes between AD and control, *GPR34* ranked 311^th^.

**Fig.5:**
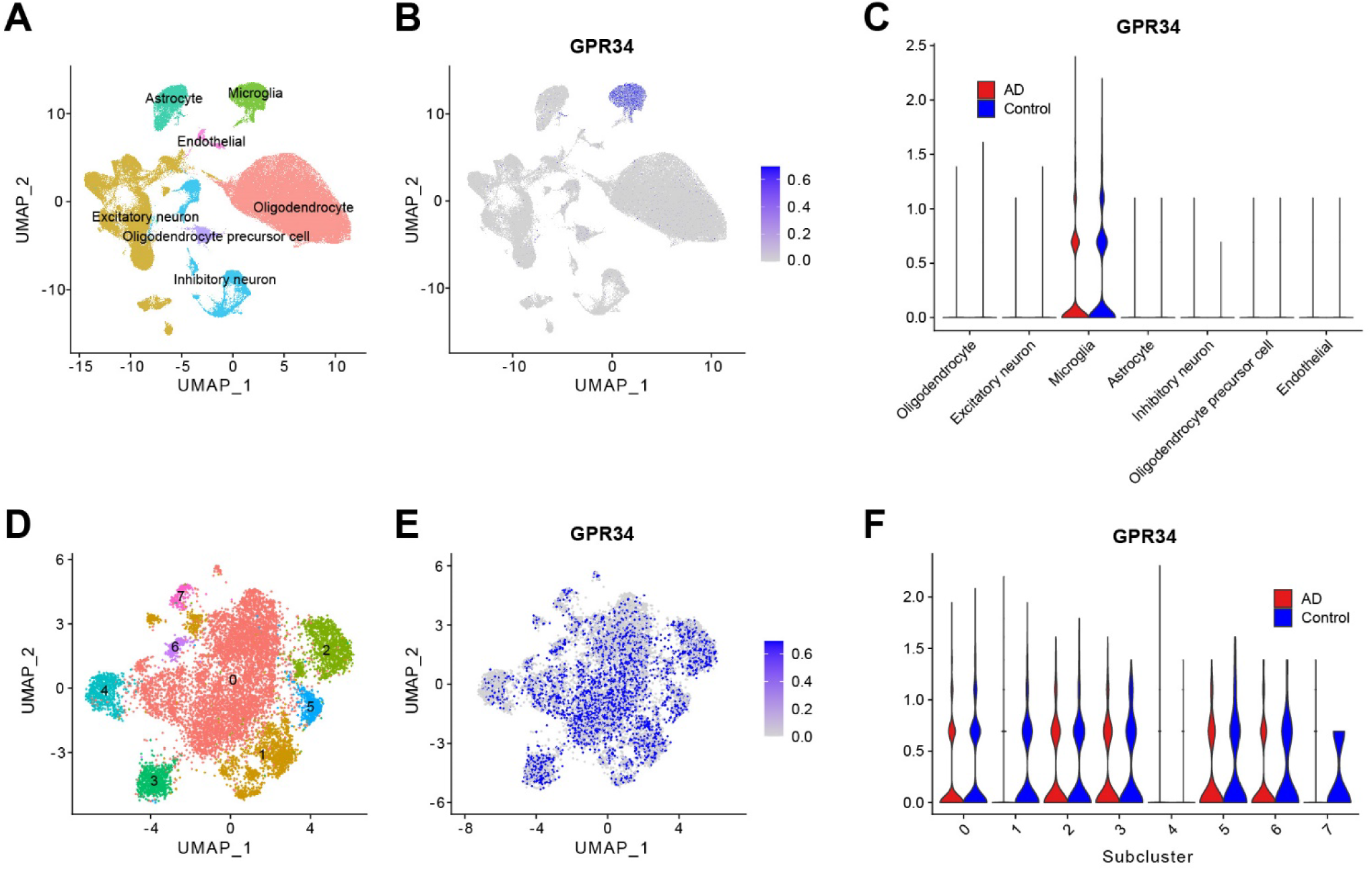
Reduced *GPR34* expression in microglia associated with Alzheimer disease ***A***–***C*** snRNA-seq analysis of the frontal cortex from 15 AD patients and 8 controls in the Japanese cohort. UMAP visualization of major cell types (***A***). Feature plots showing the expression of *GPR34* in AD and control samples (***B***). Expression levels of *GPR34* across cell types are shown as mean ± SEM (***C***). A significant reduction in *GPR34* expression was observed in AD microglia compared to controls (log_2_FC = −0.49, adjusted *p* = 0.00004). ***D-F*** Subcluster analysis of the microglial population. UMAP visualization of microglial subclusters (0-7) identified by unsupervised clustering of the microglial population from panel ***A*** (***D***). Feature plot showing the expression of GPR34 (***E***). Expression levels of *GPR34* in each microglial subcluster are shown as mean ± SEM (***F***).

To further characterize the microglial population (total 11,458 cells), we performed subcluster analysis and identified 8 distinct subclusters (0–7) (Fig. 5*D*). The feature plot of *GPR34* expression on the same UMAP space (Fig. 5*E*) suggested a relatively broad expression across the subclusters, which was further detailed in the violin plot (Fig. 5*F*). This plot revealed that *GPR34* expression was not prominently enriched in any specific subcluster, except for subcluster 4, which exhibited significantly lower expression. When comparing AD and control samples within each subcluster, a significant downregulation of *GPR34* was observed in subcluster 0 (log_2_FC = −0.51, *p* = 0.0007). Although not statistically significant, there was a trend towards decreased *GPR34* expression in the AD group across other subclusters as well. Notably, subcluster 0, comprising 6,453 cells (56% of the total) and being the largest subcluster, was characterized by the expression of microglial genes such as *CSF1R* and *P2RY12*, suggesting that it represents homeostatic microglia (*SI Appendix*, Fig. S4 *A* and *B*). On the other hand, subcluster 4, which exhibited the lowest *GPR34* expression, was enriched for macrophage markers like *CD163*, indicating its macrophage identity (*SI Appendix*, Fig. S4*C*). This finding is interesting given that *GPR34* was previously identified as specifically expressed in microglia compared to macrophages in mice (12). Our data suggest a similar specificity in humans. To further support our findings, we reanalyzed publicly available AD snRNA-seq datasets using the interactive web-based platform TACA (https://taca.lerner.ccf.org/), specifically focusing on the study by Gerrits *et al.* (32). This dataset uniquely enabled the dissection of Aβ and tau pathology contributions by including samples from the occipital cortex (OC), which exhibits Aβ pathology but minimal tau pathology in AD, and the occipitotemporal cortex (OTC) region, which harbors both Aβ and tau pathologies. This analysis revealed that in the OC region, where only Aβ accumulation occurs, GPR34 expression was significantly reduced in microglia from both AD patients and healthy controls with low-level Aβ pathology (CTR+) compared to pathology-free controls (*SI Appendix*, Table S2 and Fig. S5). Intriguingly, in the OTC region affected by both Aβ and tau pathologies, GPR34 levels were decreased in the CTR+ group with mild Aβ accumulation, but paradoxically increased in the AD group exhibiting substantial tau pathology. This suggests a potential differential effect of tau pathology on GPR34 expression, which warrants further investigation. Nonetheless, these findings corroborate our observation of reduced GPR34 expression in microglia associated with Aβ pathology. Next, we examined age-related changes in GPR34 expression using published microglial RNA-seq datasets of both human and mouse origins. In humans, while Galatro *et al.* (33) did not show significant age-dependent changes, Olah *et al.* (34) indicated a substantial decrease (log_2_FC = −1.20, adjusted *p*-value = 0.044) in the aged population compared to young individuals. In contrast, multiple mouse studies consistently revealed a significant reduction in GPR34 levels in aged microglia compared to young adults (*SI Appendix*, Fig. S6) (35–37), suggesting that an age-related decline in microglial GPR34 expression may be a conserved phenomenon across species.

The concordant findings from human AD and aging mouse models highlight a common pattern of decreased microglial GPR34 expression, which could impair Aβ clearance capability. This age-related reduction in GPR34 levels/activity may contribute to Aβ accumulation in the aging brain and exacerbate AD pathogenesis. Collectively, these results underscore the importance of maintaining or enhancing microglial GPR34 activity, for which GPR34 agonists represent a potential therapeutic strategy to promote effective Aβ clearance and prevent pathological Aβ accumulation.

## Discussion

In this study, we identified the G protein-coupled receptor GPR34 as a key regulator of microglial phagocytosis of Aβ fibrils. Activating GPR34 with a selective agonist, M1, significantly enhances the phagocytosis of Aβ fibrils in both mouse and human microglia, highlighting the translational potential of targeting GPR34 for promoting Aβ clearance. Importantly, a single intrahippocampal injection of M1 in a β-amyloidosis mouse model substantially increased microglial phagocytosis of Aβ, providing compelling evidence for the *in vivo* efficacy of GPR34 activation. Mechanistically, M1 suppresses intracellular cAMP levels, which inversely correlate with microglial Aβ phagocytosis activity, suggesting that GPR34-mediated cAMP signaling plays a crucial role in regulating Aβ clearance. Furthermore, transcriptomic analyses revealed a significant reduction in *GPR34* expression associated with AD pathology, indicating that decreased GPR34 levels may contribute to impaired Aβ clearance and disease progression. Collectively, our findings establish GPR34 as a promising therapeutic target for AD and underscore the potential of GPR34 agonists as a novel strategy to promote Aβ clearance and mitigate AD pathogenesis.

The inverse correlation between cAMP levels and fAβ phagocytosis is consistent with previous reports demonstrating the inhibitory role of cAMP in phagocytosis (26–28). These studies suggest that elevated cAMP levels can suppress phagocytic activity, possibly by modulating key signaling pathways or cytoskeletal reorganization processes involved in particle engulfment. Interestingly, the selective impact of M1/forskolin on fAβ uptake, but not on the internalization of soluble Aβ species, aligns with the notion that distinct mechanisms govern the uptake of different Aβ forms (25). On the other hand, the reason why M1 did not affect the uptake of myelin or *E. coli* particles, which are also internalized through phagocytosis, remains unclear. However, it is possible that cAMP exerts selective effects on Aβ receptors. In this regard, it is particularly intriguing that M1 failed to augment Aβ uptake in *Tyrobp*-deficient mice, which lack functional TREM2. Given the established role of TREM2 in mediating microglial phagocytosis of Aβ, it is plausible that GPR34 activation may modulate TREM2-dependent fAβ uptake by regulating cAMP signaling. However, further investigation is needed to elucidate the precise molecular mechanisms underlying this interaction. Furthermore, the Aβ fibrils used in this study form larger aggregates compared to myelin and *E. coli*, and the molecular mechanisms of phagocytosis are known to differ depending on the target size (38). Thus, these substrates may be internalized through distinct pathways with varying sensitivity to cAMP regulation.

cAMP has been implicated in the regulation of various microglial functions (39, 40). Interestingly, microglia express several Gi/o-coupled receptors, including GPR34, P2RY12, P2RY13, CX3CR1, and GPR84 (12), but the functional significance of this apparent redundancy remains elusive. While it is natural to assume that these receptors converge on the same cellular response due to the rapid diffusion of cAMP, evidence suggests that cAMP and its metabolic enzymes can be localized beneath phagocytic cups (41, 42), and cAMP can form spatially restricted signaling compartments (43). This compartmentalization raises the possibility that GPR34 and other Gi/o-coupled receptors may regulate distinct pools of cAMP with unique functional consequences, such as modulating fAβ phagocytosis. Future studies aimed at dissecting the spatial and functional organization of cAMP signaling in microglia will be crucial to understanding the specific roles of these receptors in microglial function and AD pathogenesis.

Our snRNA-seq analysis of a Japanese AD patient cohort and reanalysis of the Gerrits dataset consistently revealed a significant reduction in *GPR34* expression in AD microglia, in agreement with previously published data (32, 44). Intriguingly, the Gerrits dataset suggests that this reduction is more closely associated with Aβ accumulation than with tau pathology. In contrast, experimental studies using β-amyloidosis mouse models, such as the APP/PS1 (45, 46) and NLGF mice (47), have reported increased *Gpr34* expression in microglia. While this discrepancy between human and mouse data could be attributed to species-specific differences in GPR34 regulation, an alternative explanation may lie in the age-dependent decline of GPR34 expression observed in both human and mouse datasets. Interestingly, a similar age-associated decrease has been reported for another microglial gene, *Cd22*, which impairs phagocytic activity (35). We hypothesize that the age-related reduction in *GPR34* levels may contribute to impaired Aβ clearance and increased Aβ accumulation in the aging brain, potentially increasing the risk of developing AD. However, the further decrease in *GPR34* expression observed in AD patients compared to controls suggests that additional factors, such as genetic predisposition, may exacerbate *GPR34* downregulation in AD. The precise mechanisms underlying the downregulation of *GPR34* remain elusive; however, our findings underscore the potential significance of this process in AD pathogenesis and highlight the complex interplay between *GPR34* expression, aging, and AD pathology.

While M1 is a promising compound for promoting microglial Aβ phagocytosis, its structure suggests limited blood-brain barrier permeability, necessitating direct brain administration in this study and precluding a thorough evaluation of long-term efficacy. The development of orally bioavailable GPR34 agonists with improved brain penetrance would enable more comprehensive investigations into the therapeutic potential of GPR34 activation in AD models. In this regard, our recent elucidation of the GPR34-M1 binding structure (19) may facilitate the rational design of novel GPR34 agonists. However, it is also important to consider potential side effects related to GPR34’s roles in immune regulation and neuropathic pain (24). Future studies should carefully examine the safety and tolerability of novel GPR34 agonists in preclinical AD models.

In summary, our findings identify GPR34 as a novel therapeutic target for enhancing microglial phagocytosis of Aβ and highlight the potential of GPR34 agonists as a disease-modifying strategy for AD. Further research into the development of potent GPR34 agonists and their long-term safety and efficacy could lead to novel therapeutic approaches for this devastating disorder.

## Materials and methods

Full methods are available in *SI Appendix, Supplementary Methods*.

### Mice

All animal procedures were conducted in accordance with the protocols approved by the Institutional Animal Care and Use Committee at the University of Tokyo (UT) (protocol no. P5-04). See *SI Appendix* for further details.

### Aβ uptake assay in primary microglia

Primary microglia were prepared from neonatal C57BL/6J mice as described previously (48). For the Aβ uptake assay, cells were serum-starved in DMEM for 30 min and then treated with fluorescently labeled Aβ species (monomer, oligomer, or fibril) and either M1 or vehicle in DMEM containing 1% FBS for 3 hours. Cells were then detached, and for experiments involving Aβ oligomers or fibrils, the cell suspension was mixed with trypan blue solution to quench any extracellular fluorescence. The internalized Aβ fluorescence was quantified using flow cytometry. Data were normalized to the MFI of control cells treated with fAβ alone (i.e., without M1 or vehicle treatment). See *SI Appendix* for further details.

### Differentiation of Human Induced Pluripotent Stem Cell-derived Microglia and Aβ Uptake Assay

Microglia-like cells were differentiated from human induced pluripotent stem cells (iPSCs) using a previously described protocol (31). Briefly, iPSCs were genetically engineered to express the transcription factor *SPI1* under a doxycycline-inducible system. The differentiated microglia-like cells were used for Aβ uptake assays, where they were treated with fluorescently labeled Aβ fibrils and M1 for 4.5 hours. The internalized Aβ fluorescence was quantified using flow cytometry. See *SI Appendix* for further details.

### Stereotaxic injection of M1 and measurement of microglial uptake of Aβ *in vivo*

To assess the effect of M1 on microglial uptake of Aβ *in vivo*, 15–17-month-old male and female NLGF mice were used. Mice were anesthetized, and 0.8 µL of 250 µM M1 was injected into one hippocampus, while an equal volume of vehicle was injected into the contralateral hippocampus using stereotaxic coordinates. Nine hours after M1 injection, mice were administered methoxy-X04 intraperitoneally (MX04) to label Aβ fibrils *in vivo*. Three hours later, mice were perfused, and the hippocampi were dissected. Microglial cells were isolated and the percentage of MX04-positive microglia (CD11B+ CD45+ PI−) was assessed by flow cytometry. See *SI Appendix* for further details.

### Human samples

Autopsied human brain tissues from 15 AD patients and 8 controls without neurodegenerative pathology (Table S1) were obtained from the Brain Bank for Aging Research via the Japan Brain Bank Net. Written informed consent was obtained from the family of the deceased. Dissected brain tissues were quickly frozen with dry ice powder and stored at −80°C. Single-nucleus RNA sequencing (snRNA-seq) analysis was performed at Niigata University (NU) using frontal cortex samples. This research was approved by the Ethics Committees of the Tokyo Metropolitan Institute for Geriatrics and Gerontology (TMIG) (permit number: OF-23), NU (G2018-0034), and the UT (5–15).

### Isolation of nuclei and single-nucleus RNA sequencing

Nuclei were isolated from fresh frozen postmortem brain tissue, and for each sample, approximately 10,000 nuclei were processed for snRNA-seq using the Chromium Next GEM Single Cell 3’ Kit v3.1 (10x Genomics). Raw sequencing data were processed using the Cell Ranger pipeline (v7.0.0, 10x Genomics) for demultiplexing, barcode assignment, read alignment to the human reference genome (GRCh38-2020-A), and gene counting. Subsequent analysis was performed by using the R package *Seurat* (v5.0.1). See *SI Appendix* for further details.

### Tools and resources used in the preparation of this work

The authors employed Anthropic’s Claude for assistance in revising and refining the manuscript. After using this tool, the authors reviewed and edited the content as needed and take full responsibility for the content of the publication. Illustrations were created with BioRender.com. Chemical structures were generated using Marvin (Chemaxon).

### Statistical analysis

Statistical analyses and graphical presentations were performed using GraphPad

Prism 10. The specific statistical tests used and sample sizes for each experiment are indicated in the corresponding figure legends. P-values lower than 0.05 were considered statistically significant, except for the snRNA-seq analysis, and are represented by asterisks: *p < 0.05, **p < 0.01, ***p < 0.001, ****p < 0.0001. Non-significant differences (p > 0.05) are indicated by “ns”.

## Data, Materials, and Software Availability

The snRNA-seq data generated in this study are currently being prepared for the deposition in the DNA Data Bank of Japan (DDBJ; https://www.ddbj.nig.ac.jp/index.html). All other data are included in the manuscript and/or supporting information.

## Acknowledgments

This work was supported by the JSPS Grants-in-Aid for Scientific Research (KAKENHI) under grant numbers 19H01015, 23H00394 (to T. Tomita), 19K07080, 22K07344 (to S.T.), 21H05633 (to H.W.), 21K07271 (to A.M.), and 22H04923 (CoBiA) (to Y.S. and S.M.); AMED under grant numbers JP22dm0207072 (to T. Tomita), JP22dm0207073 (to T. Tomita and S.T.), JP21bm0804003, JP23bm1123046, JP23kk0305024 (to H.O.), JP23wm0525019 (to M.K. and T.I.), JP24dk0207060 (to T.I., A.M., and M.K.), and JP21wm0425019 (to Y.S. and S.M.); JST Moonshot R&D under grant number JPMJMS2024 (to T. Tomita); Integrated Research Initiative for Living Well with Dementia (IRIDE) of the TMIG (to Y.S. and S.M.); MHLW Research on rare and intractable diseases Program Grant Number JPMH23FC1008 (to Y.S.); ONO Medical Research Foundation (to S.T.). We would like to thank Dr. Shinya Yamanaka (Kyoto University) for the kind gifts of the 201B7 iPSC line.

## Author Contributions

Conceptualization, S.T., J.A., and T. Tomita; Methodology, S.T., W.W., H.E, I.S., H.W., H.O., N.H., A.M., M.K., and T.I.; Investigation, H.E., N.H., M.H., A.A., and J.O.; Resources, T.O., J.A., M.M., Y.S., S.M., A.M., T.I., H.W., H.O., T. S., T.C.S, and T. Takai; Writing – Original Draft, S.T. and H.E.; Review & Editing, S.T.; Project Administration, S.T., H.W. and A.M.; Supervision, S.T., T. Tomita, J.A., H.O., and I.T.

## References

1. D. J. Selkoe, J. Hardy, The amyloid hypothesis of Alzheimer’s disease at 25 years. EMBO Mol Med 8, 595–608 (2016).

2. T. Jonsson, et al., A mutation in APP protects against Alzheimer’s disease and age-related cognitive decline. Nature 488, 96–9 (2012).

3. J. Sevigny, et al., The antibody aducanumab reduces Aβ plaques in Alzheimer’s disease. Nature 537, 50–6 (2016).

4. C. H. van Dyck, et al., Lecanemab in Early Alzheimer’s Disease. N Engl J Med 388, 9–21 (2023).

5. D. V Hansen, J. E. Hanson, M. Sheng, Microglia in Alzheimer’s disease. J Cell Biol 217, 459–472 (2018).

6. S. Takatori, W. Wang, A. Iguchi, T. Tomita, Genetic Risk Factors for Alzheimer Disease: Emerging Roles of Microglia in Disease Pathomechanisms. Adv Exp Med Biol 1118, 83–116 (2019).

7. T. Jonsson, et al., Variant of TREM2 associated with the risk of Alzheimer’s disease. N Engl J Med 368, 107–16 (2013).

8. R. Guerreiro, et al., TREM2 variants in Alzheimer’s disease. N Engl J Med 368, 117–27 (2013).

9. Y. Wang, et al., TREM2 lipid sensing sustains the microglial response in an Alzheimer’s disease model. Cell 160, 1061–71 (2015).

10. T. R. Jay, et al., Disease Progression-Dependent Effects of TREM2 Deficiency in a Mouse Model of Alzheimer’s Disease. J Neurosci 37, 637–647 (2017).

11. S. Parhizkar, et al., Loss of TREM2 function increases amyloid seeding but reduces plaque-associated ApoE. Nat Neurosci 22, 191–204 (2019).

12. M. L. Bennett, et al., New tools for studying microglia in the mouse and human CNS. Proc Natl Acad Sci U S A 113, E1738–46 (2016).

13. S. Krasemann, et al., The TREM2-APOE Pathway Drives the Transcriptional Phenotype of Dysfunctional Microglia in Neurodegenerative Diseases. Immunity 47, 566–581.e9 (2017).

14. T. Sugo, et al., Identification of a lysophosphatidylserine receptor on mast cells. Biochem Biophys Res Commun 341, 1078–87 (2006).

15. A. Inoue, et al., TGFα shedding assay: an accurate and versatile method for detecting GPCR activation. Nat Methods 9, 1021–9 (2012).

16. H. Kitamura, et al., GPR34 is a receptor for lysophosphatidylserine with a fatty acid at the sn-2 position. J Biochem 151, 511–8 (2012).

17. A. Uwamizu, et al., Lysophosphatidylserine analogues differentially activate three LysoPS receptors. J Biochem 157, 151–60 (2015).

18. S. Nakamura, et al., Non-naturally Occurring Regio Isomer of Lysophosphatidylserine Exhibits Potent Agonistic Activity toward G Protein-Coupled Receptors. J Med Chem 63, 9990–10029 (2020).

19. T. Izume, et al., Structural basis for lysophosphatidylserine recognition by GPR34. Nat Commun 15, 902 (2024).

20. I. Liebscher, et al., Altered immune response in mice deficient for the G protein-coupled receptor GPR34. J Biol Chem 286, 2101–10 (2011).

21. E. Jäger, et al., Dendritic Cells Regulate GPR34 through Mitogenic Signals and Undergo Apoptosis in Its Absence. J Immunol 196, 2504–13 (2016).

22. X. Wang, et al., GPR34-mediated sensing of lysophosphatidylserine released by apoptotic neutrophils activates type 3 innate lymphoid cells to mediate tissue repair. Immunity 54, 1123–1136.e8 (2021).

23. J. Preissler, et al., Altered microglial phagocytosis in GPR34-deficient mice. Glia 63, 206–15 (2015).

24. A. Sayo, et al., GPR34 in spinal microglia exacerbates neuropathic pain in mice. J Neuroinflammation 16, 82 (2019).

25. S. Mandrekar, et al., Microglia Mediate the Clearance of Soluble Aβ through Fluid Phase Macropinocytosis. The Journal of Neuroscience 29, 4252–4262 (2009).

26. D. M. Aronoff, C. Canetti, M. Peters-Golden, Prostaglandin E2 inhibits alveolar macrophage phagocytosis through an E-prostanoid 2 receptor-mediated increase in intracellular cyclic AMP. J Immunol 173, 559–65 (2004).

27. S. A. Kalamidas, et al., cAMP synthesis and degradation by phagosomes regulate actin assembly and fusion events: consequences for mycobacteria. J Cell Sci 119, 3686–94 (2006).

28. C. H. Serezani, et al., Prostaglandin E2 suppresses bacterial killing in alveolar macrophages by inhibiting NADPH oxidase. Am J Respir Cell Mol Biol 37, 562–70 (2007).

29. T. Saito, et al., Single App knock-in mouse models of Alzheimer’s disease. Nat Neurosci 17, 661–3 (2014).

30. T. Kaifu, et al., Osteopetrosis and thalamic hypomyelinosis with synaptic degeneration in DAP12-deficient mice. J Clin Invest 111, 323–32 (2003).

31. I. Sonn, et al., Single transcription factor efficiently leads human induced pluripotent stem cells to functional microglia. Inflamm Regen 42, 20 (2022).

32. E. Gerrits, et al., Distinct amyloid-β and tau-associated microglia profiles in Alzheimer’s disease. Acta Neuropathol 141, 681–696 (2021).

33. T. F. Galatro, et al., Transcriptomic analysis of purified human cortical microglia reveals age-associated changes. Nat Neurosci 20, 1162–1171 (2017).

34. M. Olah, et al., A transcriptomic atlas of aged human microglia. Nat Commun 9, 539 (2018).

35. J. V Pluvinage, et al., CD22 blockade restores homeostatic microglial phagocytosis in ageing brains. Nature 568, 187–192 (2019).

36. A. L. Thomas, M. A. Lehn, E. M. Janssen, D. A. Hildeman, C. A. Chougnet, Naturally-aged microglia exhibit phagocytic dysfunction accompanied by gene expression changes reflective of underlying neurologic disease. Sci Rep 12, 19471 (2022).

37. S. R. Ocañas, et al., Microglial senescence contributes to female-biased neuroinflammation in the aging mouse hippocampus: implications for Alzheimer’s disease. J Neuroinflammation 20, 188 (2023).

38. D. Schlam, et al., Phosphoinositide 3-kinase enables phagocytosis of large particles by terminating actin assembly through Rac/Cdc42 GTPase-activating proteins. Nat Commun 6, 8623 (2015).

39. L.-P. Bernier, et al., Nanoscale Surveillance of the Brain by Microglia via cAMP-Regulated Filopodia. Cell Rep 27, 2895–2908.e4 (2019).

40. M. Merlini, et al., Microglial Gi-dependent dynamics regulate brain network hyperexcitability. Nat Neurosci 24, 19–23 (2021).

41. K. B. Pryzwansky, A. L. Steiner, J. K. Spitznagel, C. L. Kapoor, Compartmentalization of cyclic AMP during phagocytosis by human neutrophilic granulocytes. Science 211, 407–10 (1981).

42. M. N. Ballinger, T. Welliver, S. Straight, M. Peters-Golden, J. A. Swanson, Transient increase in cyclic AMP localized to macrophage phagosomes. PLoS One 5, e13962 (2010).

43. J. Z. Zhang, et al., Phase Separation of a PKA Regulatory Subunit Controls cAMP Compartmentation and Oncogenic Signaling. Cell 182, 1531–1544.e15 (2020).

44. S. Morabito, et al., Single-nucleus chromatin accessibility and transcriptomic characterization of Alzheimer’s disease. Nat Genet 53, 1143–1155 (2021).

45. N. Ma, et al., Whole-Transcriptome Analysis of APP/PS1 Mouse Brain and Identification of circRNA-miRNA-mRNA Networks to Investigate AD Pathogenesis. Mol Ther Nucleic Acids 18, 1049–1062 (2019).

46. L.-L. Lin, et al., GPR34 Knockdown Relieves Cognitive Deficits and Suppresses Neuroinflammation in Alzheimer’s Disease via the ERK/NF-κB Signal. Neuroscience 528, 129–139 (2023).

47. A. Sobue, et al., Microglial gene signature reveals loss of homeostatic microglia associated with neurodegeneration of Alzheimer’s disease. Acta Neuropathol Commun 9, 1 (2021).

48. Y. Kato, et al., Protocol for gene knockdown using siRNA in primary cultured neonatal murine microglia. STAR Protoc 5, 102867 (2024).

