## Supporting Information for "Selective agonism of GPR34 stimulates microglial uptake and clearance of amyloid β fibrils"

**This PDF file includes:**

Supplementary Methods

Figures S1 to S6

Tables S1 and S2

SI References

Supplementary Methods

**Mice**

*NLGF* (IMSR Cat# RBRC06344, RRID:IMSR_RBRC06344) (1) and *Tyrobp*^−/−^ (2) mice were described previously (3). Pregnant C57BL/6J mice for preparing primary microglia were purchased from SLC. All animal procedures were conducted in accordance with the protocols approved by the Institutional Animal Care and Use Committee at the University of Tokyo (protocol no. P5-04). Mice were housed under standard conditions, including a 12-hour light/dark cycle, a temperature of 22°C, and humidity ranging from 40 to 60%. They were provided with *ad libitum* access to food and water. Both male and female mice were used for the analyses.

**Preparation of Aβ species**

For Aβ monomer preparation, HiLyte Fluor 647-labeled human Aβ_1-40_ (Anaspec) was dissolved in dimethyl sulfoxide (DMSO) to obtain a stock concentration of 1 mg/mL. This stock solution was stored at −80°C and, prior to use, diluted in culture medium to a final concentration of 1 µg/mL.

For Aβ oligomer and fibril preparation, stock solutions of HiLyte Fluor 488-labeled human Aβ_1-42_ (Anaspec) and unlabeled human Aβ_1-42_ (Peptide Institute) were prepared using 1,1,1,3,3,3-Hexafluoro-2-propanol (HFIP) and stored at −30°C until use. Before use, the labeled and unlabeled Aβ peptides were mixed at a molar ratio of 1:20. The HFIP was subsequently evaporated using a Speed Vac. The Aβ peptides were then reconstituted in 10 µL of DMSO per 50 µg of peptide, followed by dilution in 490 µL HBSS(+), yielding a 22 µM solution. After passing through a 0.22-µm PVDF filter, the solution was incubated overnight at 4°C (for oligomers) or 37°C (for fibrils).

For pHrodo conjugation of Aβ fibrils, a 22 µM solution of unlabeled human Aβ_1-42_ peptides was fibrillized as described above. Post-fibrilization, the fibril solution was mixed with pHrodo Green STP Ester (Thermo Fisher Scientific) at a molar ratio of 1:50 (Aβ:pHrodo dye). After 30 min of incubation at room temperature, the fibrils were pelleted by centrifugation at 20,000 × g for 10 min. The supernatant was removed, and the pellet was washed with HBSS(+). This washing step was repeated once, and the fibrils were finally resuspended in HBSS(+).

**Preparation of M1**

M1 was synthesized as described previously (4–6). A series of 100× concentrated stock solutions of M1 were prepared in Dulbecco’s phosphate-buffered saline (D-PBS) containing 0.1% (w/v) protease- and fatty-acid-free bovine serum albumin (BSA; SERVA Cat#11945) and stored at −30°C until use. For *in vitro* experiments, the stock solutions were diluted 100-fold in the appropriate medium to achieve the desired final concentrations.

**Aβ uptake assay in mouse primary microglia**

Primary microglia were prepared from C57BL/6J mice as described previously (3, 7). For Aβ uptake assays, microglial cells were seeded at a density of 5 × 10^4^ cells per well in a volume of 200 µL of 10% L929-conditioned medium (LCM) in 48-well tissue-culture-treated plates and cultured for 24 hours at 37°C. Before initiating the assay, cells were serum-starved in DMEM for 30 min. The medium was then replaced with DMEM containing 1% FBS (DMEM/FBS) and Aβ species (1 µg/mL for monomer, 1 µM for oligomer, and 2 µM for fibril), with either M1 or vehicle. After a 3-hour treatment, cells were detached by incubation with Dulbecco’s phosphate-buffered saline (D-PBS) supplemented with 0.125% trypsin and 1 µM ethylenediaminetetraacetic acid for 5 min at 37°C. The enzymatic rection was terminated by adding D-PBS containing 2% FBS. The cells were then collected by gentle pipetting, passed through a 40-µm cell strainer, and kept on ice until analysis. For experiments involving Aβ oligomers or fibrils, the cell suspension was mixed with an equal volume of 250 µg/mL trypan blue to quench any extracellular fluorescence. The fluorescence derived from internalized Aβ was measured using an MA900 flow cytometer (SONY).

For siRNA knockdown experiments, primary microglial cells were seeded in 12-well low-adherence plates at a density of 1.5 × 10^5^ cells per well in 10% LCM. After 48 hours, the cells were transfected with siRNA using Lipofectamine RNAiMAX (Thermo Fisher Scientific) according to the manufacturer’s instructions. Twenty-four hours post-transfection, the culture medium was replaced with fresh medium, and cells were incubated for an additional 48 hours before being subjected to Aβ uptake and qPCR analyses.

**Quantitative real-time polymerase chain reaction**

Total RNA was isolated using ISOGEN (NIPPON GENE) and stored at −80°C until use. For complementary DNA (cDNA) synthesis, the isolated RNA was reverse-transcribed using ReverTra Ace qPCR RT Master Mix with gDNA Remover (TOYOBO). The resultant cDNA was then diluted tenfold with Milli-Q water. Quantitative real-time PCR assays were performed in duplicate on a LightCycler 480 Instrument II (Roche) using THUNDERBIRD SYBR qPCR Mix (TOYOBO) and the following primer sets: *Gpr34* (forward: 5'-GCAGCTTCTGAAGTATGGCCT-3', reverse: 5'-AAGCCAGCTGTCAACTGAAGTA-3') and *Gapdh* (forward: 5'-AGGTCGGTGTGAACGGATTTG-3', reverse: 5'-TGTAGACCATGTAGTTGAGGTCA-3'). To generate standard curves, serial dilutions of the samples were prepared at 1/5, 1/25, and 1/125 using Milli-Q water. Relative gene expression changes were assessed using the ΔCT method and normalized to *Gapdh* expression.

***Escherichia coli* BioParticles uptake assay**

Primary microglial cells were incubated with Alexa Fluor 488-conjugated *Escherichia coli* K-12 BioParticles (Invitrogen) at a concentration of 20 µg/mL in DMEM/FBS for 20 min. Following the incubation, cells were washed with D-PBS and then detached using 0.125% trypsin and 1 µM EDTA for 5 min at 37°C. The enzymatic reaction was terminated by adding D-PBS containing 2% FBS. Cells were subsequently collected by gentle pipetting and passed through a 40-µm cell strainer. The collected cells were kept on ice until fluorescence measurement. To quench any extracellular fluorescence, an equal volume of 250 µg/mL trypan blue was added to the cell suspension immediately before analysis. The internalized fluorescence (488 nm) was quantified using a flow cytometer.

**Myelin debris preparation and uptake assay**

Myelin debris was prepared from the brains of C57BL/6J mice following the protocol described by Rolfe *et al.* (8) The myelin stock solution was adjusted to a protein concentration of 100 mg/mL in DPBS and stored at −80°C until use. For the assay, the myelin stock was quickly thawed and resuspended using a 29-gauge needle syringe. It was then mixed with pHrodo Green STP Ester reagent (1 mg/mL) at a volume ratio of 10:1. This mixture was incubated at room temperature in the dark for 30 min. Subsequently, pHrodo-labeled myelin was isolated by centrifugation at 20,000 × g for 10 min, discarding the supernatant and resuspending the pellet in D-PBS.

For the uptake assay, cells were treated with pHrodo-labeled myelin at a concentration of 1 mg/mL in DMEM/FBS for 3 h. Following the incubation, cells were trypsinized and resuspended in a solution containing 20 mM CHES-NaOH (pH 9.0) and 150 mM NaCl. The internalized fluorescence (488 nm) was measured using a flow cytometer.

**Cyclic AMP concentration measurement**

Primary microglial cells were seeded in 48-well low-adherence plates at 2.5 × 10^4^ cells/well in 100 µL of 10% LCM. After 24 hours of culturing, the cells were rinsed twice with HBSS(+) and subsequently incubated in HBSS(+) for 15 min at room temperature. The cells were then treated with 20 µL of HBSS(+) containing 100 µM forskolin (dissolved in DMSO) and/or 10 µM M1, followed by another 15-minute incubation at room temperature. The cells were lysed, and cAMP concentrations were determined using the cAMP-Glo Assay (Promega) according to the manufacturer's protocol.

**Cell lysis and immunoblotting**

Primary microglial cells were rinsed with ice-cold PBS, lysed and sonicated in RIPA buffer containing protease and phosphatase inhibitors. The protein concentration of the samples was determined using the bicinchoninic acid protein assay (TaKaRa). 5× SDS-PAGE sample buffer and 2-mercaptoethanol (to a final concentration of 1% (v/v)) were added to the samples, which were then heated for 5 min at 100°C.

Proteins were separated by Tris-glycine SDS polyacrylamide gel electrophoresis and transferred to PVDF membranes. The membranes were blocked for 1 h at room temperature with 3% BSA in TBS containing 0.1% (v/v) Tween 20 (TBST), followed by three washes with TBST. Membranes were incubated with the appropriate dilutions of primary antibodies, including pan-Akt (clone C67E7; Cat# 4691), phospho-Akt (Ser473) (clone D9E; Cat# 4060), and pan-ERK1/2 (clone 137F5; Cat# 4695) from Cell Signaling Technology and p-ERK (clone E-4; Cat# sc-7383) from Santa Cruz Biotechnology, in TBST containing NaN_3_ overnight at 4°C. After three washes with TBST, membranes were incubated with the appropriate peroxidase-conjugated secondary antibody (1:2,000) in TBST for 1 h at room temperature. After three washes with TBST, membranes were incubated with Immunostar luminescence working solution. Signals were detected using an ImageQuant LAS 4000 (GE Healthcare) and processed using Image J/Fiji (1.53c).

**Culture of human induced pluripotent stem cells (iPSCs)**

The human iPSC lines 201B7 (9) and RPC802 (ReproCell Inc, cat# RCRP002N) were cultured in a feeder-free condition. All iPSCs were maintained in 6-well plates (Corning) coated with iMatrix-511. iPSCs were cultured in StemFit/AK02N (Ajinomoto) medium supplemented with penicillin/streptomycin in a humidified incubator (37°C, 5% CO_2_), and the medium was changed every other day. The genetically engineered iPSCs expressing *SPI1* in a doxycycline-dependent manner were described previously (10). Human ethics approval was obtained from the Ethics Committee in Keio University School of Medicine (Approval Number: 20080016).

**Differentiation of human induced pluripotent stem cell-derived microglia and Aβ uptake assay**

Microglia derived from human induced pluripotent stem cells (hiMGLs) were generated as described previously (10). Once differentiated, hiMGLs were seeded on poly-D-lysine-coated 48-well plates at a density of 7.5 × 10^4^ cells per well. Twenty-four hours after seeding, a half volume of the culture medium was freshly added. Following an additional 24-hour incubation, the cells were treated with 2 µM Aβ fibrils and either M1 or vehicle for 4.5 hours. The cells were then detached using Accutase at 37°C for 5 min. This enzymatic reaction was terminated by adding D-PBS. The cells were subsequently collected by gentle pipetting and passed through a 40-µm cell strainer. The internalized Aβ fluorescence was quantified as described above.

**Stereotaxic injection of M1 and measurement of microglial uptake of Aβ *in vivo***

To assess the effect of M1 on microglial uptake of Aβ in vivo, male and female NLGF mice aged 15–17 months were used. Mice were anesthetized with a subcutaneous injection of a mixture containing 0.3 mg/kg medetomidine, 4.0 mg/kg midazolam, and 5.0 mg/kg butorphanol. After exposing the skull, 0.8 µL of 250 µM M1 was injected into one hippocampus, while an equal volume of vehicle was injected into the contralateral hippocampus. The injections were performed using the following stereotaxic coordinates: A/P: −1.7 mm, M/L: +/−1.6 mm, D/V: −1.9 mm from bregma. Before injection, the syringe needle was first inserted 0.1 mm deeper than the target to create a small pocket. The injection was performed at a flow rate of 0.2 µL/min. After the injection, the needle was left in place for an additional 2–3 min before being carefully retracted. The surgical incision was sutured, and mice were revived by a subcutaneous injection of 3.0 mg/kg atipamezole. Throughout the procedure, the mice’s body temperature was maintained using a heating pad. Post-surgery, mice had ad libitum access to food and water.

Twelve hours after M1 injection, microglial cells were isolated from the hippocampi to assess Aβ uptake. To fluorescently label Aβ fibrils in vivo, 10 mg/kg MX04 was administered intraperitoneally to the mice three hours before analysis. Mice were then anesthetized, perfused with a minimum of 20 mL ice-cold PBS, and the hippocampi were removed. A single-cell suspension was obtained using the GentleMACS Octo dissociator with heaters and the Adult Brain Dissociation Kit (Miltenyi). Subsequently, myelin debris was removed using the Debris removal solution (Miltenyi). After blocking Fc receptors with 0.5 µg/mL of CD16/32 antibody (Miltenyi #130-092-374) for at least 10 min on ice, the cell suspension was incubated with Alexa Fluor 647 anti-mouse CD45 antibody (BioLegend #103124) and REAfinity™ anti-mouse CD11B antibody (Miltenyi #130-113-805) for another 10 min on ice. After centrifugation, cells were resuspended and analyzed by flow cytometry. Two to three minutes before measurement, 1 µg/mL propidium iodide was added to stain dead cells. CD11B+ CD45+ PI− cells were gated, and the percentage of MX04-positive cells was assessed.

**Isolation of nuclei from frozen brain tissue**

We isolated nuclei from approximately 20 mg of fresh frozen postmortem brain tissue using the Minute Single Nucleus Isolation Kit for Neuronal Tissues/Cells (Invent Biotechnologies) according to the manufacturer’s protocol. Isolated nuclei were stained with ReadyCount Blue Nuclear Stain (Thermo Fisher Scientific) and counted on the Countess II FL automated cell counter (Thermo Fisher Scientific).

**Single-nucleus RNA sequencing**

Isolated nuclei were processed for single-nucleus RNA sequencing using the Chromium Next GEM Single Cell 3' Kit v3.1 (10x Genomics) following the manufacturer’s protocol. Briefly, for each library, approximately 10,000 nuclei were partitioned into gel beads-in-emulsion on the Chromium Controller instrument (10x Genomics). Reverse transcription and cDNA amplification were performed, and Illumina-compatible adapter sequences were added to the final libraries via PCR. The libraries were sequenced on Illumina’s NextSeq2000 instrument. Raw sequencing data were processed using the Cell Ranger pipeline (v7.0.0, 10x Genomics) for demultiplexing, barcode assignment, read alignment to the human reference genome (GRCh38-2020-A), and gene counting. Subsequent analysis followed the vignettes from the R package *Seurat* (v5.0.1). Initially, Cell Ranger output data were imported into R (v4.3.2) via the *SoupX* (v1.6.2) package to eliminate ambient RNA contamination. Nuclei were excluded if the number of detected features (genes) was less than 200 or greater than 14,000 and the percentage of mitochondrial genes was greater than 5%. Count data were normalized using the *SCTransform* v2 function while adjusting for the proportion of mitochondrial genes, and then multiplets were removed using the *DoubletFinder* (v2.0.4) package. The *RunPCA* function (npcs = 30) was run, and all datasets were integrated using the *Harmony* (v1.2.0) package through the *IntegrateLayers* function. The functions *FindNeighbors* (dims = 30), *FindClusters* (resolution = 0.2), and *RunUMAP* (dims = 1:30) were performed, and clusters were visualized through the *DimPlot* function. To identify gene expression patterns of major cell type-specific markers, we used the *FindAllMarkers* function (min.pct = 0.25, logfc.threshold = 0.25, test.use = “MAST”), adjusting for the difference in diagnosis (AD or control) with the latent.vars option. We then annotated the cell types according to the expression patterns of the following genes: *NRGN*, *SLC17A7*, and *CAMK2A* for excitatory neuron, *GAD1* and *GAD2* for inhibitory neuron, *AQP4* and *GFAP* for astrocyte, *MBP*, *MOBP* and *PLP1* for oligodendrocyte, *CSF1R*, *CD74* and *C3* for microglia, *VCAN*, *PDGFRA* and *CSPG4* for oligodendrocyte precursor cell, and *FLT1* and *CLDN5* for endothelial cell. The microglia cluster was subjected to subclustering, and the same procedure as described above was conducted. Two rounds of integration steps were repeated to remove clusters contaminated with non-microglial markers, and the cleaned microglia dataset was divided into eight clusters through *FindClusters* (resolution = 0.1). To identify differentially expressed genes between AD and control, we used the *FindMarkers* function (min.pct = 0.10, logfc.threshold = 0.1, test.use = "MAST") while adjusting for sex and *APOE* genotypes. *P*-values were adjusted using the Bonferroni correction for multiple testing, and *p*-values less than 0.01 were considered statistically significant.

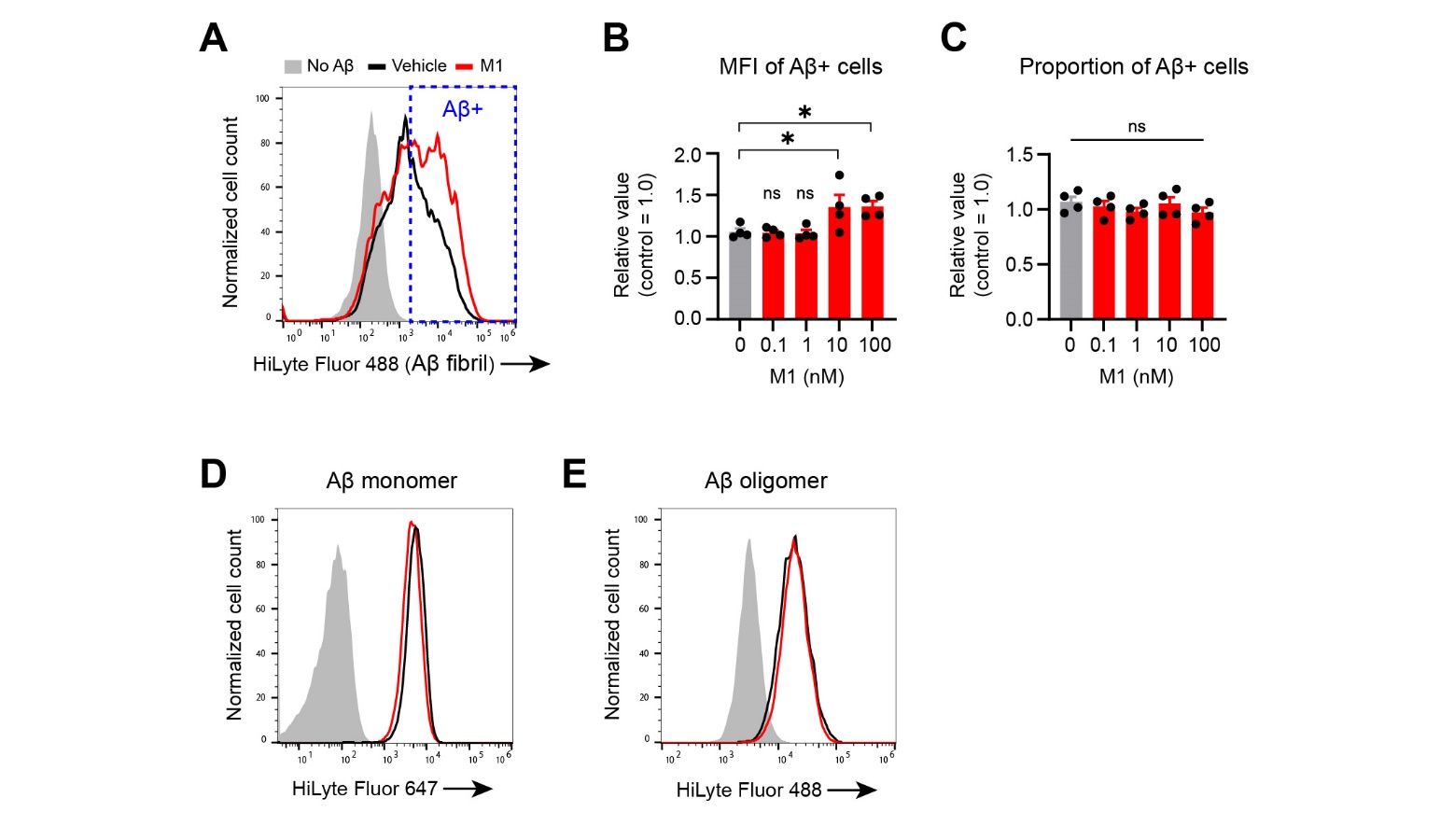

Fig. S1. M1 effects on the MFI and proportion of Aβ+ cells (monomer, oligomer, fibril)

***A***–***C*** Representative flow cytometry plots (***A***) and quantification of the mean fluorescence intensity (MFI) (***B***) or proportion (***C***) of fibrillar Aβ+ cells, showing the effect of M1 treatment. The gated population used for quantification is highlighted with a blue dashed box in ***A***. Data in ***A*** and ***B*** are the same as those presented in Fig. 1 *B* and *C*, respectively. Data are normalized to the values of the control cells (N=4). Bars represent the mean ± SEM. Dunnett’s test.

***D***, ***E*** Representative flow cytometry plots showing the effect of M1 on the uptake of fluorescently labeled Aβ monomers (***D***) and oligomers (***E***) by primary microglia. See Fig. 1*G* for quantification.

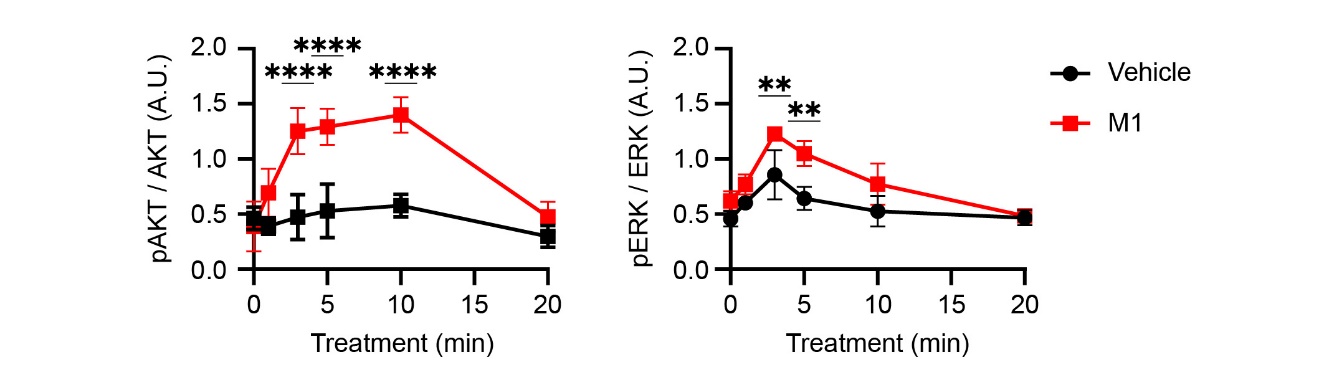

Fig. S2. M1 effects on the phosphorylation of AKT and ERK in primary microglia

Quantification of phosphorylated AKT (p-AKT) and ERK (p-ERK) levels in primary microglia treated with 10 nM M1 for the indicated times. The p-AKT and p-ERK levels were normalized to total AKT and ERK levels, respectively. Data are presented as mean ± SD (N=3). Two-way ANOVA followed by Šídák’s multiple comparisons test.

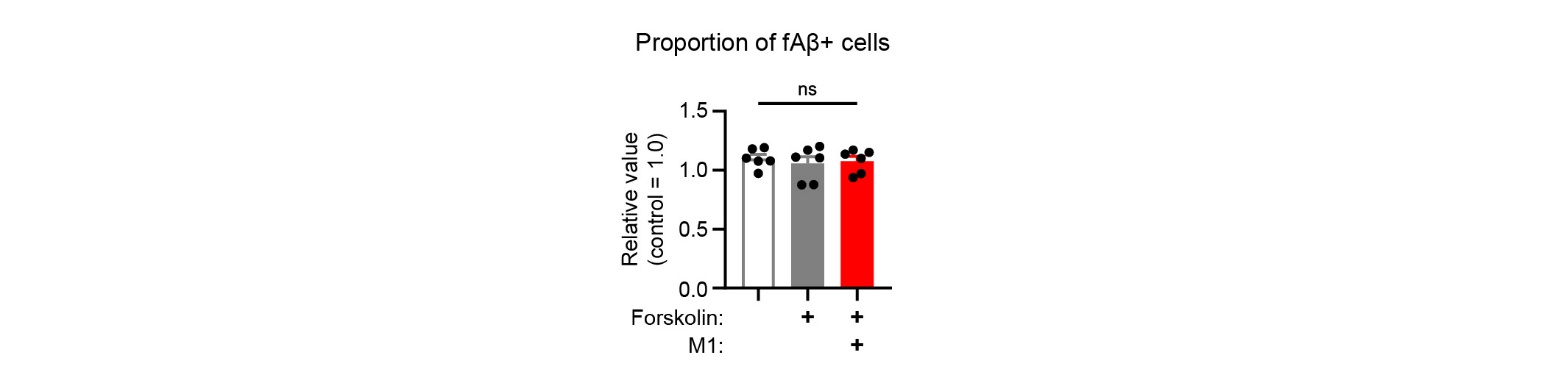

Fig. S3. Effects of forskolin and M1 on the proportion of fibrillar Aβ-positive cells

Quantification of the proportion of fibrillar Aβ (fAβ)+ cells upon treatment with vehicle, forskolin, or forskolin + M1. Data were derived from the same flow cytometry analysis as in Fig. 2*C*, but here the proportion of fAβ+ cells is shown. The proportion was normalized to the values of the control cells. Data are presented as mean ± SEM (N=6). One-way repeated measures ANOVA followed by Tukey's multiple comparisons test.

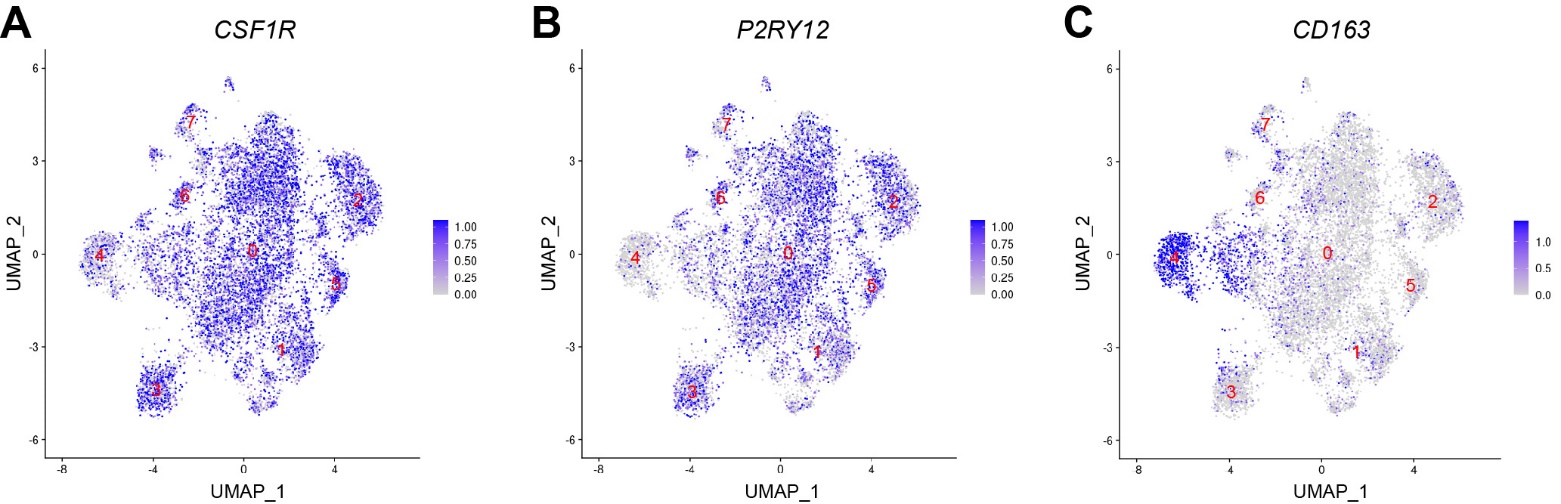

Fig. S4. Expression of myeloid marker genes across subclusters

Feature plots showing the expression levels of myeloid marker genes on the same UMAP space as in Fig. 5*D*. The microglial subclusters are indicated by the numbers in red. (***A***) Expression of *CSF1R*, a marker for homeostatic microglia. (***B***) *P2RY12*, another homeostatic microglial marker, showing prominent expression in subcluster 0. (***C***) *CD163*, a marker for macrophages, is specifically expressed in subcluster 4, indicating a non-microglial identity of this subcluster.

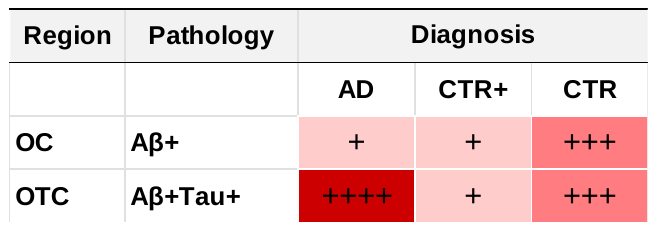

Fig. S5. Matrix summarizing the relative changes in *GPR34* expression levels in microglia from the Gerrits *et al*. dataset, related to Table S2

The matrix shows the relative expression levels of GPR34 in microglia from the occipital cortex (OC) and occipitotemporal cortex (OTC) of control (CTR), control with Aβ pathology (CTR+), and AD patients. The relative magnitude of *GPR34* expression levels is represented by the number of plus signs and the intensity of the red color within each cell. The data were obtained by reanalyzing the snRNA-seq dataset from Gerrits *et al*., 2021 (11). The corresponding fold-changes and statistical significance are provided in Table S2.

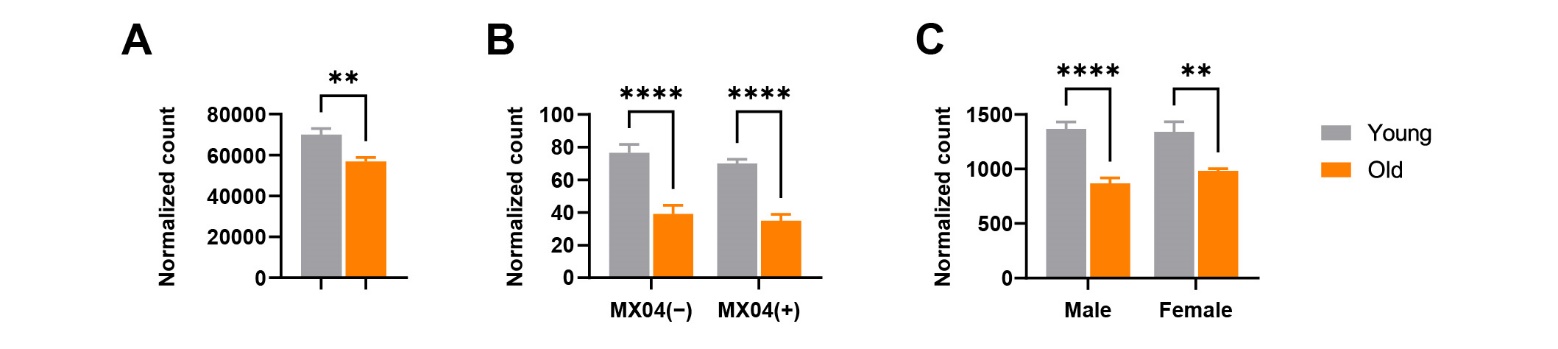

Fig. S6. Age-dependent decrease in microglial *Gpr34* expression in mouse models.

(***A***) *Gpr34* expression in microglia from young (3-month-old) and aged (20-month-old) mouse hippocampi. Data were derived from GSE127542 (12). N = 6 per group.

(***B***) *Gpr34* expression in microglia from young (< 3-month-old) and aged (> 18-month-old) mice, further separated based on their Aβ uptake status (MX04− or MX04+) determined by flow cytometry. Data were derived from GSE205803 (13). N = 5 per group.

(***C***) *Gpr34* expression in MACS-isolated microglia from young (5–6-month-old) and aged (22–25-month-old) mice, stratified by sex. Data were derived from GSE233400 (14). N = 3–5 per group.

Data are presented as mean ± SEM. Statistical significance was determined using an unpaired t-test (***A***), two-way ANOVA followed by Šídák’s multiple comparisons test (***B*** and ***C***).

Table S1. Demographic and clinical information of the Japanese snRNA-seq cohort

| ID | Diagnosis | Age | Sex | APOE |
| --- | --- | --- | --- | --- |
| AD01 | AD | 70 | Male | 4*4 |
| AD02 | AD | 74 | Male | 3*4 |
| AD03 | AD | 75 | Male | 4*4 |
| AD04 | AD | 78 | Male | 4*4 |
| AD05 | AD | 82 | Female | 3*3 |
| AD06 | AD | 82 | Female | 4*4 |
| AD07 | AD | 83 | Female | 3*4 |
| AD08 | AD | 85 | Male | 2*3 |
| AD09 | AD | 86 | Female | 4*4 |
| AD10 | AD | 87 | Male | 3*4 |
| AD11 | AD | 88 | Male | 3*4 |
| AD12 | AD | 88 | Male | 4*4 |
| AD13 | AD | 92 | Male | 3*3 |
| AD14 | AD | 93 | Male | 3*3 |
| AD15 | AD | 98 | Female | 2*3 |
| CT01 | Control | 57 | Male | 3*4 |
| CT02 | Control | 64 | Male | 3*3 |
| CT03 | Control | 70 | Male | 3*3 |
| CT04 | Control | 70 | Male | 3*4 |
| CT05 | Control | 73 | Female | 3*4 |
| CT06 | Control | 77 | Male | 3*3 |
| CT07 | Control | 79 | Male | 3*3 |
| CT08 | Control | 82 | Female | 3*3 |

*Diagnosis*: pathological diagnosis (AD or normal control); *Age*: age at death; *APOE*: *APOE* genotype of the participant.

Table S2. Differential expression analysis of *GPR34* in microglia from the Gerrits *et al*. dataset.

| **Gene** | **Brain region** | **Pathology** | **Comparison** | **Cell type** | **log2FC** | **FDR** |
| --- | --- | --- | --- | --- | --- | --- |
| *GPR34* | OC | Aβ+ | AD vs CTR | Microglia | **-0.662** | **2.0E-28** |
| *GPR34* | OC | Aβ+ | CTR+ vs CTR | Microglia | **-0.178** | **2.0E-04** |
| *GPR34* | OC | Aβ+ | AD vs CTR+ | Microglia | 0.116 | 1.0E+00 |
| *GPR34* | OTC | Aβ+Tau+ | AD vs CTR | Microglia | **0.038** | **8.0E-06** |
| *GPR34* | OTC | Aβ+Tau+ | CTR+ vs CTR | Microglia | **-0.216** | **4.0E-12** |
| *GPR34* | OTC | Aβ+Tau+ | AD vs CTR+ | Microglia | **0.254** | **5.0E-10** |

Differential expression analysis of *GPR34* in microglia from the occipital cortex (OC) and occipitotemporal cortex (OTC) of control (CTR), control with Aβ pathology (CTR+), and AD patients. The data were obtained by reanalyzing the snRNA-seq dataset from Gerrits *et al*., 2021 (11). Positive log2 fold-change (FC) values indicate upregulation in the comparison group relative to the reference group. FDR: false discovery rate-adjusted *P*-value.

**SI References**

1.	T. Saito, *et al.*, Single App knock-in mouse models of Alzheimer’s disease. *Nat Neurosci* **17**, 661–3 (2014).

2.	T. Kaifu, *et al.*, Osteopetrosis and thalamic hypomyelinosis with synaptic degeneration in DAP12-deficient mice. *J Clin Invest* **111**, 323–32 (2003).

3.	A. Iguchi, *et al.*, INPP5D modulates TREM2 loss-of-function phenotypes in a β-amyloidosis mouse model. *iScience* **26**, 106375 (2023).

4.	M. Ikubo, *et al.*, Structure-activity relationships of lysophosphatidylserine analogs as agonists of G-protein-coupled receptors GPR34, P2Y10, and GPR174. *J Med Chem* **58**, 4204–19 (2015).

5.	M. Sayama, *et al.*, Probing the Hydrophobic Binding Pocket of G-Protein-Coupled Lysophosphatidylserine Receptor GPR34/LPS1 by Docking-Aided Structure-Activity Analysis. *J Med Chem* **60**, 6384–6399 (2017).

6.	S. Nakamura, *et al.*, Non-naturally Occurring Regio Isomer of Lysophosphatidylserine Exhibits Potent Agonistic Activity toward G Protein-Coupled Receptors. *J Med Chem* **63**, 9990–10029 (2020).

7.	Y. Kato, *et al.*, Protocol for gene knockdown using siRNA in primary cultured neonatal murine microglia. *STAR Protoc* **5**, 102867 (2024).

8.	A. J. Rolfe, D. B. Bosco, E. N. Broussard, Y. Ren, In Vitro Phagocytosis of Myelin Debris by Bone Marrow-Derived Macrophages. *J Vis Exp* **2017** (2017).

9.	K. Takahashi, *et al.*, Induction of pluripotent stem cells from adult human fibroblasts by defined factors. *Cell* **131**, 861–872 (2007).

10.	I. Sonn, *et al.*, Single transcription factor efficiently leads human induced pluripotent stem cells to functional microglia. *Inflamm Regen* **42**, 20 (2022).

11.	E. Gerrits, *et al.*, Distinct amyloid-β and tau-associated microglia profiles in Alzheimer’s disease. *Acta Neuropathol* **141**, 681–696 (2021).

12.	J. V Pluvinage, *et al.*, CD22 blockade restores homeostatic microglial phagocytosis in ageing brains. *Nature* **568**, 187–192 (2019).

13.	A. L. Thomas, M. A. Lehn, E. M. Janssen, D. A. Hildeman, C. A. Chougnet, Naturally-aged microglia exhibit phagocytic dysfunction accompanied by gene expression changes reflective of underlying neurologic disease. *Sci Rep* **12**, 19471 (2022).

14.	S. R. Ocañas, *et al.*, Microglial senescence contributes to female-biased neuroinflammation in the aging mouse hippocampus: implications for Alzheimer’s disease. *J Neuroinflammation* **20**, 188 (2023).
